# Lightning-caused disturbance frequency and severity varies with topography in an Afromontane forest

**DOI:** 10.1101/2025.05.13.653675

**Authors:** Evan M. Gora, Alain Senghor K. Ngute, Rodrigue Batumike, Aida Cuni-Sanchez, Gerard Imani, Beth A. Kaplin, Drew Bantlin, Robert Bitariho, Bianca Zoletto, Nikolaos Petridis, Douglas Sheil, Martin J.P. Sullivan

## Abstract

Lightning is an important agent of tropical tree mortality, but we know little about lightning-caused disturbances beyond one lowland tropical forest in Panama. Here we quantified variation in the frequency and severity of visually-detectable lightning-caused disturbances across topography in the montane forests of Nyungwe National Park, Rwanda. This was the first systematic assessment of lightning-caused disturbance in Africa and a preliminary test of the hypothesis that forest tolerance to lightning is higher where lightning disturbances are more common. Lightning-caused disturbances were observed six-times more frequently on ridges than in valleys, but lightning-caused disturbances in valleys tended to cause more tree-level damage than those on ridges. Overall, lightning disturbances tended to kill a lower proportion of trees in this Afromontane forest than in previously documented lightning disturbances in tropical America. We also observed less tree-level damage to *Syzygium* spp.(Myrtaceae) compared to the community-wide average, providing support for taxonomic differences in lightning tolerance. Our results indicate that lightning disturbance severity differs within and among sites, mediated in part by differences in lightning tolerance among tree species, and provide some support for the hypothesis that disturbance severity declines with disturbance frequency.

## Introduction

Tropical forests cover less than 10% of the global land area yet store approximately 50% of terrestrial vegetation carbon (Pan *et al*., 2024) and contribute more than one-third of primary productivity (Beer *et al*., 2010). The tropical forest carbon sink has weakened in recent decades as tree biomass mortality increased (Brienen *et al*., 2015), but African forests appear to be more resilient to these changes (Hubau *et al*., 2020; Bennett *et al*., 2021). Problematically, we know very little about the proximate drivers of tree mortality, much less why patterns of tree mortality are changing or why the changes differ among regions (Hubau *et al*., 2020; Sullivan *et al*., 2020; McDowell *et al*., 2018). In particular, our understanding of storm-driven tree mortality and associated carbon fluxes lags far behind our understanding of other climate-sensitive agents of tree mortality, like drought and fire (McDowell *et al*., 2018; Brando *et al*., 2012; Gora and Esquivel-Muelbert, 2021).

Lightning strikes are a major, climate-sensitive component of storm-driven mortality. Lightning causes 40-50% of mortality of large trees (>60 cm in diameter) and 16% of biomass mortality in a lowland forest in Panama (Gora *et al*., 2021; Yanoviak *et al*., 2020), which is the only place where lightning has been systematically quantified in a tropical forest. Moreover, lightning strikes tropical forests 35-67 million times each year and forests that experience more lightning strikes have fewer large trees, higher rates of biomass turnover, and less aboveground biomass (Gora *et al*., 2020a), suggesting that lightning is a major driver of tree mortality and carbon dynamics pantropically. Lightning frequency is increasing (Harel and Price, 2020; Lavigne, Liu and Liu, 2019; Raghavendra *et al*., 2018), and its contributions to changes in the tropical forest carbon sink will likely be influenced by the ability of forests to acclimate to increased lightning frequency.

Lightning-caused disturbance is particularly likely to be important in Afromontane forests, which function as key carbon stores (Cuni-Sanchez *et al*., 2021) and sinks (Cuni-Sanchez *et al*., 2024). Specifically, the montane forests of the Albertine Rift experience some of the highest lightning frequencies in the world (Kigotsi, Soula and Georgis, 2018; Soula *et al*., 2016; Gora *et al*., 2020a) and in Afromontane forests, trees above 50 cm diameter, which are more likely to be struck by lightning (Gora *et al*., 2020b), account for ∼60% of aboveground forest biomass (Cuni-Sanchez *et al*., 2021). However, we know virtually nothing about the role of lightning in Afromontane forests. Given evidence that lightning frequency is increasing ca. 7% per decade in this region (Harel and Price, 2020), understanding the contemporary contributions of lightning to forest turnover is critical to understanding the future of these forests and their carbon dynamics.

Field observations of lightning strikes describe remarkably similar patterns of lightning damage across tropical American and Indomalayan forests. Typically, lightning appears to attach to a canopy tree and then electric current flows aboveground across branches or air gaps (i.e., “flashover”) to secondarily damage many neighboring trees (Anderson, 1964; Furtado, 1935; Magnusson, Lima and De Lima, 1996; Gora and Yanoviak, 2020). The only system for systematically studying lightning strikes, which used both electrical and optical sensors in central Panama, showed that an average strike damages 23.6 trees, kills 5.3 of these damaged trees, and causes 7.36 Mg of aboveground tree biomass turnover across an area of 451 m^2^ (Gora *et al*., 2021). By contrast, to our knowledge, the only published descriptions of lightning strikes in African tropical forests (1) documented a single tree that briefly caught on fire in Gabon (Caroline, Lee and Mackanga-Missandzou, 1996) and (2) putative lightning scars described in Uganda (Zoletto *et al*., 2023). Systematic field data are therefore needed to understand lightning-caused disturbance in African forests.

In montane forests, lightning strikes are expected to be concentrated on exposed ridges. At a fine scale (<15 m), taller trees are more likely to be struck than their shorter neighbors, essentially intercepting strikes that would otherwise hit neighbors if they had the same height (Gora *et al*., 2020b). Basic physics principles indicate that the distance of this “protection” effect is approximately equal to the difference in height between a focal tree and its neighbors (Uman, 2008). Across flat terrain this effect is limited. However, in the topographically complex terrain common to montane forests, the difference in heights between laterally proximate trees can be substantial. Accordingly, exposed trees on ridges likely “protect” trees across expansive downslope areas. This expectation is supported by limited observations in North American temperate forests (Yanoviak *et al*., 2015), but the concentration of lightning disturbances on ridges has not been systematically tested.

Variation in lightning frequency could produce differences in forest-level lightning tolerance. Tree species vary in their tolerance to lightning (Richards *et al*., 2022), and therefore lightning disturbances could filter out lightning-intolerant trees or change forest structure such that forests experiencing high lightning frequency, such as mountain ridges, have more lightning-tolerant tree species than forests that rarely experience lightning. Alternatively, it is possible that very few trees can survive lightning strikes regardless of their frequency, leading to limited effects on community composition and forest-level lightning tolerance. If we can understand how the frequency of lightning disturbance is related to forest-level lightning tolerance, then that will help us understand both contemporary disturbance regimes and future forest responses to changing lightning frequency.

To begin understanding the effects of lightning in African forests, here we used visual detection methods developed and tested in tropical America (Gora and Yanoviak, 2020) to quantify the frequency and severity of lightning-caused disturbances across topography in the Afromontane forests of Nyungwe National Park in Rwanda. We hypothesize that (1) lightning disturbances are more common on ridges than valleys, (2) lightning disturbances are less severe where they are more common (i.e., ridges) and (3) the characteristics of lightning disturbances in this African forest are similar to those observed on other continents. We examine these hypotheses using field observations of tree and forest patch-scale lightning disturbance effects.

## Methods

### Study area

The study was conducted in Nyungwe National Park, which covers 1013 km^2^. Elevation ranges from 1600 to 2950 m a.s.l. The region has a bimodal rainfall regime with long rains in March–May and short rains in September–December (Sun *et al*., 1996). At Uwinka research station (2,465 m asl) at the center of the park, annual rainfall is 1900 mm and average daytime and nighttime temperatures are 15.7 °C and 13.5 °C, respectively. The park contains grasslands, wetlands, bamboo-dominated forest, mixed-species old-growth and secondary montane forests, much of the latter created by past human disturbance. The most abundant tree species in old-growth forests are *Syzygium guineense, Carapa grandiflora, Ocotea usambarensis, Faurea saligna*, and *Ficalhoa laurifolia*, while secondary forests are dominated by *Macaranga kilimandscharica* (Nyirambangutse et al. 2017).

Although Nyungwe National Park was only established in March 2004, it was first gazetted as a forest reserve in the early 1930s (Masozera, 2002). This park is an area recognized for its high biodiversity and endemism (Plumptre *et al*., 2002) with at least 1105 species of plants, 280 species of birds, and 86 species of mammals (Plumptre *et al*., 2002).

### Trail surveys and lightning identification

We used transect surveys to locate lightning-caused disturbances from diagnostic patterns of lightning damage. Specifically, flashover damage is diagnostic of lightning (Gora *et al*., 2021; Yanoviak *et al*., 2017; Furtado, 1935), and it is visible as the defoliation of the two nearest branches of neighboring trees in a directionally biased pattern (i.e., often only the terminal ends of these branches die). Using a predefined procedure, we walked 23.29 km of existing trails while inspecting visible canopy trees for signs of flashover damage. We used a handheld GPS to record the 3-dimensional path of these trail surveys, and we categorized each section of this path as one of three types of topography (ridges, valleys, slopes) relevant to lightning strike likelihood. Specifically, we recorded ridges as the flat and rounded sections near topographic highpoints, valleys as local depressions where topography had limited influence on the shape of the upper surface of the forest canopy within the range of possible flashover (ca. 40 m laterally), and slopes as all areas in between.

If an observed tree exhibited potential flashover damage connected to two or more neighbors (i.e., ≥3 trees with potential flashover damage), then it was selected for more detailed surveys. In these detailed surveys, we visually followed the branches of all neighboring trees towards the initially identified tree to determine if the ends of these branches exhibited signs of flashover damage. For all trees that exhibited possible flashover damage, we visually inspected their neighbors using the same process. We only considered potential flashover events across <2 m of open air because the voltage needed to induce flashover is proportional to distance and flashover events >2m are rarely observed (Gora *et al*., 2021; Yanoviak *et al*., 2017; Furtado, 1935). Additionally, we excluded damaged trees that did not exhibit evidence of flashover damage to limit misclassification of incidental crown damage (e.g. from wind). The presumptive directly struck tree was identified as the canopy tree at the center of observed flashover damage.

There is uncertainty in assigning a proximate agent of disturbance. Accordingly, we performed quantitative and qualitative assessments of whether each damage event was associated with lightning: (1) we recorded the total number of flashover events visible at each site as a quantitative measure of evidence for lightning damage, and (2) we qualitatively differentiated disturbances that we were confident were caused by lightning (hereafter “high-confidence lightning disturbances”) from others where a strike event was likely, but the evidence was more limited (“low-confidence lightning disturbances”, high and low-confidence lightning disturbances collectively called “potential lightning disturbances”). High-confidence lightning disturbances were those for which flashover was apparent among nearly all trees within range to experience flashover damage (i.e., <1m in 3-dimensional space) and these patterns had not yet been obscured by decomposition. These methods were based on protocols tested in Peru (Gora and Yanoviak, 2020) after being developed in Panama, which is the only place globally where a remote lightning location system was used to reliably locate field-recorded strikes and develop these field diagnostics for lightning damage (Gora *et al*., 2021; Yanoviak *et al*., 2017). We refer to these events as “lightning disturbances” because they represent variation among lightning strikes that cause visible disturbances, but they do not include the effects of lightning strikes that do not cause disturbance. This is similar to typical methods for investigating other types of disturbance; for example, studies of wind-caused disturbance focus only on wind events that cause damage and fire disturbance research is typically restricted to fires that spread beyond their initial ignition.

### Lightning disturbance data collection

We recorded the topography (ridge, slope and valley, see above) and forest structure in the immediate vicinity of the disturbance (closed and open, with the latter being where trees were sufficiently spaced to reduce opportunities for flashover). We classified all damaged and dead trees into four diameter-based size classes (<10cm, 10-30cm, 30-50cm, and >50cm), using diameter tapes to measure trees where we were uncertain of size-class classification.

We took more detailed measurements on the size and condition of damaged trees >50cm in diameter at breast height (DBH) because trees of this size contributed 85% of total woody biomass turnover by lightning in a Panamanian forest (Gora *et al*., 2021). Specifically, we recorded tree DBH, height, genus identity, and crown condition. Crown damage severity, which is predictive of future mortality (Arellano *et al*., 2019), was evaluated using two approaches: (1) *crown loss* was estimated as the percent of branch volume lost based on broken branches and grouped into five categories: <5%, 5-25%, 25-50%, 50-75%, 75-95%, and >95% loss, and (2) *crown dieback* was estimated as the percent of leaf volume lost from existing branches in 5% increments, following previous work by (Gora *et al*., 2021). We calculated tree-level damage severity as the estimated crown loss, expressed as a proportion of the total crown, plus the product of crown dieback and 1-crown loss.

### Statistical analysis

We used a simulation approach to test if the number of lightning disturbances observed in each topographic position differed from expectation if they were randomly distributed across the length of trail sampled. Specifically, we simulated the locations of 23 (total number of high-confidence disturbances) or 44 (total number of high and low-confidence disturbances) disturbances, and we repeated this process for 10,000 simulations. The probability that each simulated lightning-caused disturbance was in each topographic position was equal to the proportion of the total trail length sampled in each habitat (i.e., 0.213 on ridges, 0.641 on slopes, and 0.135 in valleys). We estimated the probability that our observations could arise at random as the proportion of simulations that had equal or fewer lightning-caused disturbances as we observed in valleys and equal or more disturbances as observed on ridges (i.e. equivalent to one-tailed P values). It is possible that the area surveyed per-length of trail differed between habitats (if disturbances were more detectable in some settings that others), but we confirmed that the distance from trails to the observed lightning disturbances did not differ among topography classes (Kruskal-Wallis test, X^2^ = 3.6, df=2, P=0.165; mean distance from lightning disturbances to trail = 11.6m, maximum = 42m), and also repeated the simulation analysis using only lightning disturbances within 10m of the trail (i.e. where we were highly confident of detecting all disturbances, Fig. S1).

We used generalized linear models with negative binomial errors and a log link function to test whether the number of trees damaged or killed per lightning-caused disturbance differed with topographic position (ridge, slope, valley) and forest openness (open forest with gaps between trees, or closed-canopy). A negative binomial error distribution was chosen to account for overdispersion in the data. To assess mortality risk independently of disturbance size, we also modelled the proportion of affected trees that had died as a function of the same variables using a binomial generalized linear model. Here, each lightning disturbance represented a trial, with the number of trees killed representing the number of successes and the number of trees damaged but not killed representing the number of failures. Models were run using data from all disturbances, and also restricting data to high-confidence lightning disturbances.

We used models with similar error structures to test whether the per-disturbance effects of lightning (number of damaged trees, number of dead trees, proportion of affected trees that died) differed between Nyungwe and previously published studies in tropical American forests that used flashover to diagnose lightning damage (Yanoviak *et al*., 2020; Gora and Yanoviak, 2020). Specifically, these models only included forest site as a predictor with different levels for Nyungwe and tropical Americas. To estimate disturbance characteristics for the Americas, we aggregated data from Barro Colorado Island (BCI) in Panama (Yanoviak *et al*., 2020) and northeastern Peru (Gora and Yanoviak, 2020). The forests of BCI experience high lightning frequency (12.6 strikes km^-2^ yr^-1^) and their lightning strikes were primarily located using optical or electrical sensors (Yanoviak *et al*., 2017; Gora *et al*., 2021), whereas the Peruvian forest experienced below-average lightning frequency (2.1 strikes km^-2^ yr^-1^) and those lightning-caused disturbances from Peru were located using visual identification from drone imagery with on-the-ground validation using the same diagnostic methods (Gora and Yanoviak, 2020).

We also evaluated how the effects of lightning varied among individual trees. Using only large trees from the lightning disturbances (trees >50 cm DBH with N = 30 trees in high-confidence lightning disturbances and N = 42 trees across all disturbances, note this includes both presumed focal and secondarily affected trees, and that some lightning disturbances lacked trees >50cm DBH), we evaluated how the severity of tree damage differed with topography, forest openness, and among tree taxa. Both presumed focal and secondarily affected trees are included due to the uncertainty of identifying which tree was initially struck. Crown damage severity was logit transformed and included as a continuous response variable. Topography and forest openness had three and two levels respectively, as defined above. Tree taxa was a categorical variable with two levels, *Syzygium* or other genera, because *Syzygium* was the only genus with enough replicates for analysis (Fig. S2). These variables were fixed effects in a mixed effects model implemented using the lme4 R package (Bates *et al*., 2015), with a unique ID term for each disturbance as a random effect. Statistical significance of terms was established by parametric bootstrap implemented in the performance R package. Not all focal trees could be confidently identified to species (as lightning damage or mortality meant they had no leaves), so we repeated the tree-level analysis including only trees that could be identified to genus level.

## Results

### Damage characteristics of high-confidence and low-confidence lightning-caused disturbances

We located a total of 44 potential lightning-caused disturbances, of which 23 appeared high-confidence based on our protocols. Although our qualitative assessment was based on the patterns of flashover damage rather than the absolute number of events, our qualitative assessment was related to quantitative metrics, with an average of 10.2 flashover events (SD = 6.5) for high-confidence lightning disturbances and 3.9 flashover events (SD = 1.6) for lower confidence disturbances. Confidence in assigning lightning as a cause of disturbance differed between closed and open forest (60% of potential lightning disturbances in closed-canopy forest were considered high-confidence, compared to 46% in open forest), likely due to the greater opportunity for flashover to near neighbors in closed forest.

### Strike position across topography

Lightning-caused disturbances were more likely to be observed on ridges than would be expected if lightning disturbances were equally distributed across the lengths of all trails (Fig. 1). The 23.29 km of trail sampling consisted of 21.3% on ridges, 65.1% on slopes, and 13.5% in valleys, while we found 19 of 44 potential lightning disturbances on ridges (43.2%; the probability of this number of lightning disturbances or higher randomly occurring on ridges if they were randomly distributed with respect to topography is = 0.001), 23 (52.3%, P = 0.055) on slopes and only 2 of 44 in valleys (4.5%, P = 0.056). Patterns were similar, but no longer statistically significant, among high-confidence lightning disturbances: 8 of 23 were located on ridges (34.8%, P = 0.1010), 13 (56.5%, P =0.142) in slopes and 2 of 23 were in valleys (8.7%, P = 0.375). Results were similar when restricting to disturbances within 10m of trails, with significantly more lightning disturbances than expected in ridges when including both high and low confidence disturbances (Fig. S1). Lightning disturbances on slopes also appeared more frequent for trees growing in exposed positions; 8 out of 13 high-confidence and 5 out of 10 low-confidence lightning disturbances on slopes affected focal trees growing in exposed positions over steep slopes.

**Figure. 1.**
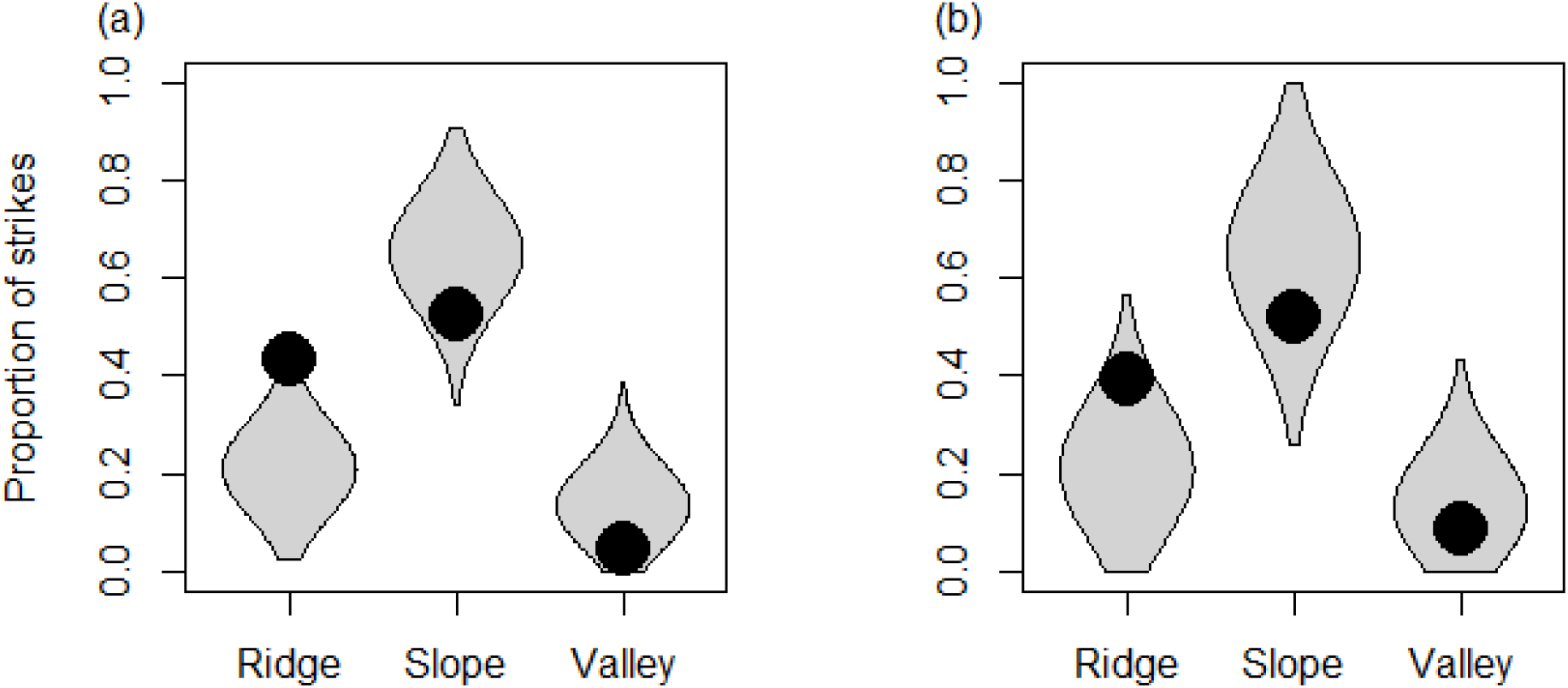
Variation in the frequency of lightning disturbances by topographic position in relation to expectations if lightning disturbances were randomly distributed with respect to topography. Points show the proportion of observed (a) potential and high-confidence strikes and (b) high-confidence strikes in each topographic category, Violin plots show the expected distribution of strike proportions from 10000 simulations where lightning disturbances were randomly assigned across topographic categories, weighted by the distance of trails surveyed in each topographic category.

### Variation in lightning disturbance characteristics

Each detected lightning disturbance caused visible damage (including death) to 10.0 ± 7.3 SD trees (Fig 2a, 13.5±8.6 for high-confidence lightning disturbances), and killed 1.8±2.1 trees (2.3±2.6 for high-confidence disturbances). Lightning disturbances killed a higher proportion of trees in valleys compared to ridges and slopes (32% of affected trees killed in valleys, compared to 15% in ridges and 16% in slopes, Fig. 2a), however, with only two disturbances in valleys, this difference was not statistically significant (binomial GLM, β_valley_vs_ridge_ = 0.794 ± 0.531 SE, z = 1.5, df = 40, P = 0.135, see Table S1 for full model results). The number of trees affected by lightning disturbances did not vary with topography (9.8 ± 8.7SD trees damaged on ridges, 10.1 ± 6.4 on slopes, 9.5 ± 4.9 in valleys), but fewer trees were damaged or killed in open forest compared to closed canopy forest (negative binomial GLMs, damage: β = -0.48 ± 0.18 SE, z = 2.7, df = 40, P = 0.006, dead: β = -0.65 ± 0.32, z = 2.0, df = 40, P = 0.044, Fig. 2a). Patterns were similar when the analysis was restricted to high confidence strikes, although differences between open and closed forest were no longer statistically significant (Table S1). Our results thus raise the potential for lighting to kill a greater proportion of trees in valleys than in ridges, but the rarity of lightning disturbances in valleys challenges statistical assessment.

**Figure 2.**
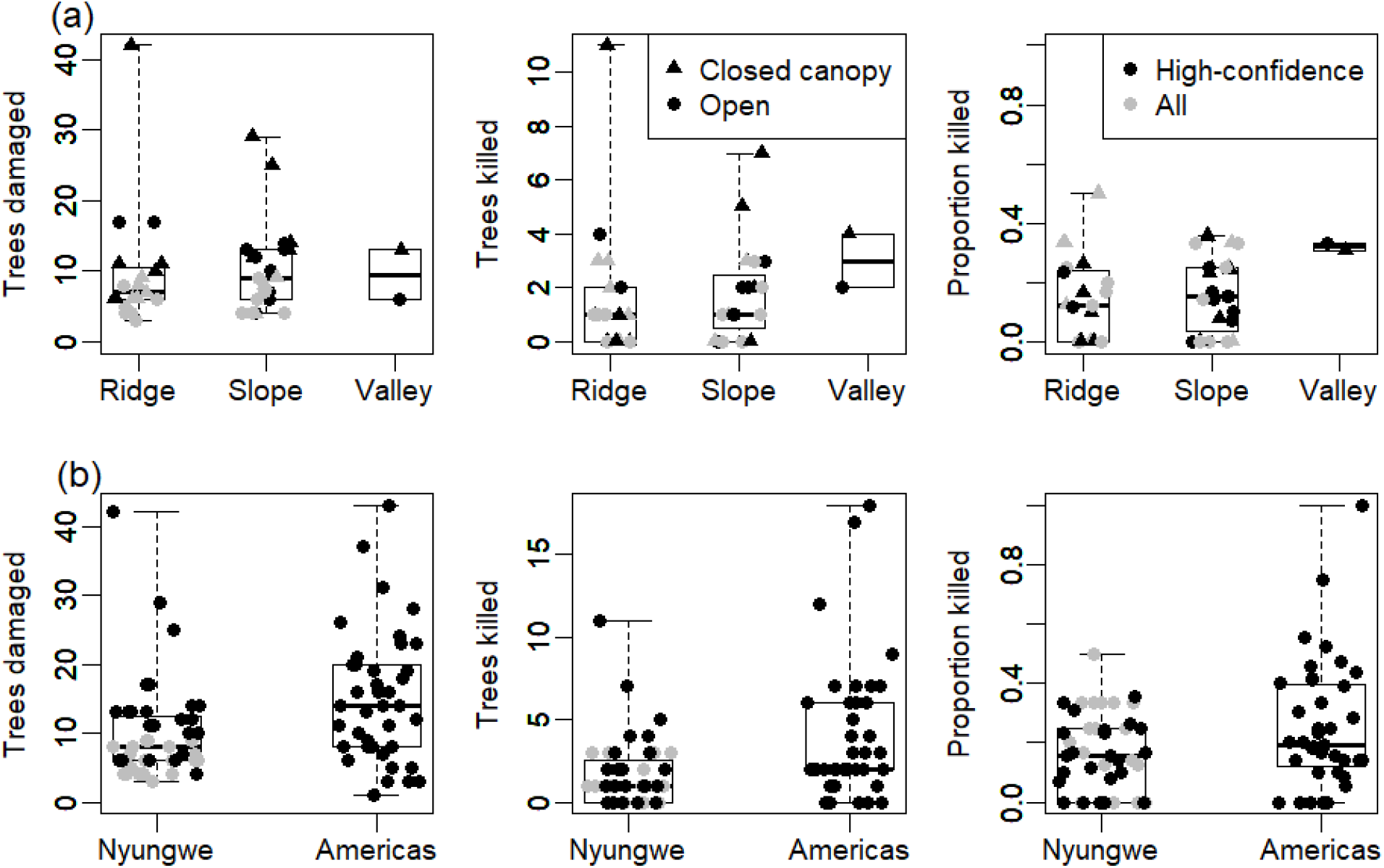
Variation in lightning disturbance characteristics (a) within Nyungwe forest across topographic positions and forest canopy closure categories and (b) between Nyungwe and previously published data from tropical America (Barro Coloardo Island, n = 32 and north-east Peru, n = 7). Black symbols indicate high-confidence lightning disturbances, and grey symbols indicate low-confidence lightning disturbances. In (a), triangles indicate disturbances in closed canopy forest, and circles disturbances in open forests.

Comparing the patterns of lightning disturbance recorded in this montane forest to the two American lowland forest sites (Panama, and northeastern Peru), lighting damaged (Negative binomial GLM, β = -0.42±0.14 SE, z = 3.1, df = 81, P = 0.002) and killed (negative binomial GLM, β = -0.83±0.23, z = 3.62, df = 81, P < 0.001) significantly fewer trees in Nyungwe than in the tropical American sites (analysis using all lightning disturbances at Nyungwe). The lower number of trees killed per strike remained marginally statistically significant when the analysis was restricted to high-confidence disturbances at Nyungwe (negative binomial GLM, β = - 0.56±0.27, z = 2.0, df = 60, P = 0.0416), but the difference in the number of damaged trees was no longer statistically significant (negative binomial GLM, β = -0.11±0.16, z = 0.7, df=60, P = 0.483). The proportion of affected trees that were killed was lower at Nyungwe (binomial GLM, all disturbances: β = -0.53±0.16, z = 3.4, df = 81, P <0.001; high-confidence disturbances: β = - 0.57±0.18, z = 3.2, df = 60, P = 0.001). Our results thus indicate that lightning-caused disturbances kill fewer trees per lightning disturbance in Nyungwe than previous observations in Panama and Peru.

### Variation in tree-level damage

*Syzygium* spp. trees exhibit much less damage than the community-wide average (difference in logit-transformed crown damage to large trees [trees >50 cm in diameter] within all lightning-caused disturbances = -2.03, 95% CI = -3.26 to -0.83, P <0.001, Fig. 3), but there were no significant differences in crown damage with topography or forest openness (Table S2). *Syzygium* made up 56% of the large trees in ridgetop lightning disturbances (n=18 trees), 48% of large trees in lightning disturbances on slopes (n = 21 trees) but none of the four large lightning-damaged trees in valleys were *Syzygium*. The lower crown damage to *Syzygium* remained when restricting the analysis to high confidence lightning disturbances, or removing trees that could not be identified to genus level (Table S2). When only high confidence lightning disturbances were included, there were significant differences in crown damage with topography (similar in ridges and slopes, but greater damage in valleys compared to ridges, difference in logit crown damage = 2.3, 95% CI = 0.26 to 4.34, P=0.026, Table S2) and forest type (greater in open forest than closed canopy forest, difference = 1.75, 95% CI = 0.16 to 3.49, P = 0.018, Table S2).

**Figure 3.**
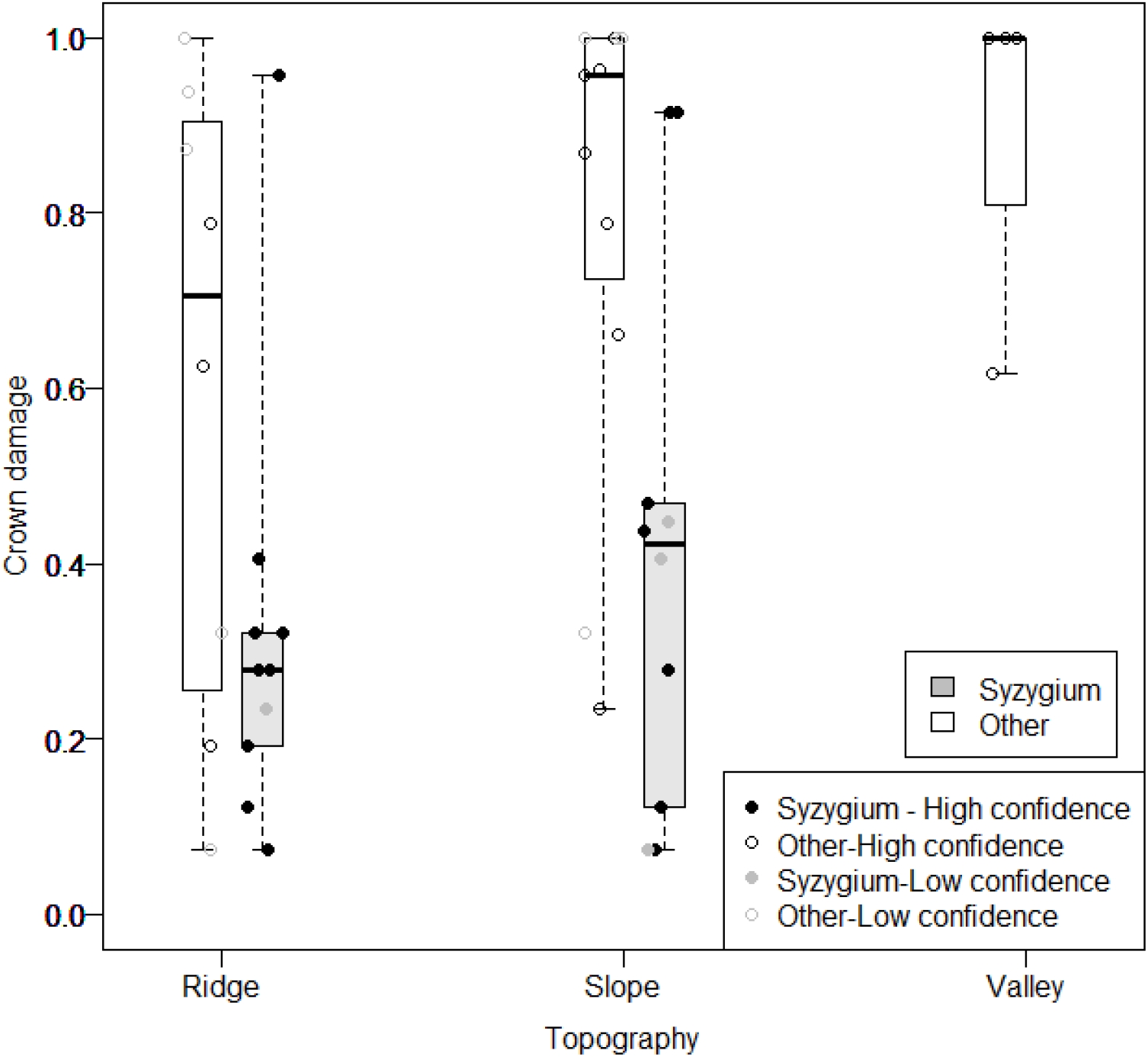
Variation in tree-level damage caused by lighting. Points show our composite crown damage metric for each large tree (>50cm DBH) in high-confidence (black symbols) and possible (grey symbols), separating out strikes on Syzygium (filled symbols, gray boxes) to other taxa (open symbols, white boxes). Boxplots have been drawn for each taxonomy-topography combination. Crown damage for each genus is shown in Fig. S2.

## Discussion

Here we provide the first systematic quantification of lightning-caused disturbance in any African tropical forest and in any montane forest pantropically. We show that lightning-caused disturbances are much more likely to be observed on ridges than in valleys, providing the best evidence to-date supporting the long-standing hypothesis that lightning strikes are more common on ridges than in valleys (reviewed by (Yanoviak *et al*., 2015)). We also provide some evidence that lightning-caused disturbances are more severe on valleys than in ridges, albeit with statistical analysis limited by the rarity of lightning disturbances in valleys. Together, these patterns are consistent with the hypothesis that forests experiencing higher lightning strike rates are less vulnerable to lightning disturbance. Overall, our data suggest that lightning is an important agent of disturbance that likely influences the composition and dynamics of tropical montane forests and African forests, but these data have limitations and there are many opportunities for future research.

The non-random concentration of lightning-caused disturbances on forest ridges has implications for landscape-scale patterns of forest structure and composition. The tendency of ridgetops to have shorter trees and lower biomass is often attributed to edaphic and hydrological differences (Detto *et al*., 2013), cloud-immersion (Fahey, Sherman and Tanner, 2016) or to wind-caused mortality (Jackson *et al*., 2021), but our data suggest that lightning-caused disturbances also contribute to these structural differences. Moreover, interspecific variation in lightning tolerance (Richards *et al*., 2022) and evidence that trees of certain species even benefit from being struck by lightning (Gora et al. in press) strongly suggest that higher lightning frequency on ridges could influence the composition of these communities. Consequently, it is possible that exposed ridges in areas of high lightning frequency, such as the Albertine Rift (Gora *et al*., 2020a), could represent a specialized habitat niche that can be exploited by tree species relatively more tolerant to lightning impacts.

If, as our results suggest, lightning disturbance severity is less severe where lightning is most frequent, this implies that forests will differ in their responses to climate-driven increases in lightning frequency (Harel and Price, 2020). The contributions of the genus *Szyzgium* to these patterns in our data are consistent with compositional shifts towards more lightning-tolerant taxa being key to forest-level tolerance (Richards *et al*., 2022). Indeed, *Syzygium guineense* (Engl.) Mildbr. is one of the most abundant (in terms of stem density) late-successional trees in Nyungwe (Nyirambangutse *et al*., 2017) and across the high lightning frequency forests of Kibira National Park in Burundi (Hakizimana, Huynen and Hambuckers, 2016) and Kahuzi-Biega National Park and Itombwe Mountains in eastern DRC (Imani *et al*., 2021; Imani *et al*., 2017). The low sample size of other genera meant we were only able to compare *Syzygium* spp. (assumed to be mainly *S. guineense* as this is locally the most abundant species in the genus) to all other taxa combined. However, it is likely that lightning tolerance varies amongst the other genera (Fig. S2). In addition to compositional shifts, our observations suggest that the open forest structure of ridgetop forests also reduce lightning disturbance severity by decreasing the number of neighboring trees with sufficient proximity to experience flashover damage. Regardless of the specific mechanism, incorporating these compensatory responses of forests to shifting disturbance regimes and increasing lightning frequency (Harel and Price, 2020; Lavigne, Liu and Liu, 2019; Raghavendra *et al*., 2018) will be key to accurately forecasting forest composition, structure, and function (Pugh *et al*., 2020).

The Albertine Rift and eastern Congo Basin forests experience exceptionally high lightning strike frequencies (Cecil, Buechler and Blakeslee, 2014; Gora *et al*., 2020a), and we expect that higher lightning strike rates could result in regional and biogeographic patterns in tolerance to lightning. The lesser severity of a typical lightning-caused disturbance in Nyungwe is consistent with this hypothesis, particularly when compared to the higher severity of lightning strikes surveyed post-hoc in Amazonian forests (Fig. 2) and in South-East Asia (Sarawak: three documented strikes killed 35.3 ± 16.6 SD trees per strike,(Anderson, 1964)). However, this difference could also arise from the biased sampling of ridges in this study focused on montane forests, which are unlikely to be representative of African forests in general, given that most tropical forests in Africa are lowland-not montane. Overall, methodological differences between studies complicates cross-continent comparisons, and explicitly testing for regional differences in lightning responses will require systematically locating and quantifying lightning strikes across biogeographic realms and elevation.

These results provide rare empirical support for long-standing hypotheses that are grounded in physical and ecological theory (reviewed by Yanoviak *et al*., 2015), but the underlying data have limitations. First, we provide surveys of visually apparent lightning disturbances, but because these data omit visually inapparent lightning strikes, such as those that do not cause meaningful damage, they represent the typical effects of lightning-caused disturbances rather than representing the per-strike or landscape-level contributions of lightning to forest dynamics. This limitation is common to the disturbance ecology literature – we often ignore weak winds that do not cause damage, fires that do not spread after their ignition, and short-term droughts that do not cause meaningful drought stress – but it is notable nonetheless. Second, it is possible that unquantified biases affected observations of lightning disturbance across topography or between open and closed-canopy forest types. The non-random location of the trails could influence the forests encountered during these surveys. Additionally, the denser forest structure of valleys relative to ridgetop forests could both (1) increase detection efficiency of lightning disturbance by providing more opportunities for flashover damage and (2) obscure other lightning-caused disturbances by reducing visibility from the trail. The competing nature of these potential biases, the large difference in disturbance density, and the strong theoretical basis for topographic effects on lightning distributions all suggest that these biases have limited effect, but we fundamentally cannot test their influence beyond confirming that patterns remain when restricting sampling to very close to trails. To overcome these limitations, it would be necessary to establish a lightning detection system to provide unbiased detections of strikes with subsequent quantification (Yanoviak *et al*., 2017).

Our results and these limitations highlight several major avenues for future research. First, these are the first systematic data describing lightning-caused disturbance in Africa, but they are unlikely to broadly represent African forests. Additional research and replication are needed to quantify the effects of lightning across many African forests, especially given its status as a lightning hotspot (Cecil, Buechler and Blakeslee, 2014; Gora *et al*., 2020a), and to confirm the differences in lightning disturbance severity between valleys and ridges. Second, we show that ridges experience higher rates of lightning disturbance, but a detailed study of lightning strike probabilities (Uman, 2008) with empirical validation is needed to properly quantify and model these effects (Gora *et al*., 2020b). Third, we provide preliminary evidence that higher lightning frequency is associated with greater forest-level lightning tolerance, but additional research is needed to uncover the mechanisms underlying *Szyzgium spp*. tolerance and to quantify the contributions of both forest structure and tree species composition to forest-level tolerance. We expect future research to find that lightning is a key agent of disturbance across many African forests with implications for forest ecology and evolution.

## Acknowledgements

Nyungwe National Park is managed by African Parks, a non-profit conservation organisation that manages protected areas on behalf of governments across Africa, through a public-private partnership with the Rwanda Development Board. Permission to conduct research was provided by Nyungwe Management Company, the Rwanda Development Board, and the Republic of Rwanda National Council for Science and Technology. We are indebted to the Nyungwe National Park rangers for their support with the research. This research was funded by Natural Environment Research Council grant NE/W003872/1.

## Supporting Information

**Figure S1.**
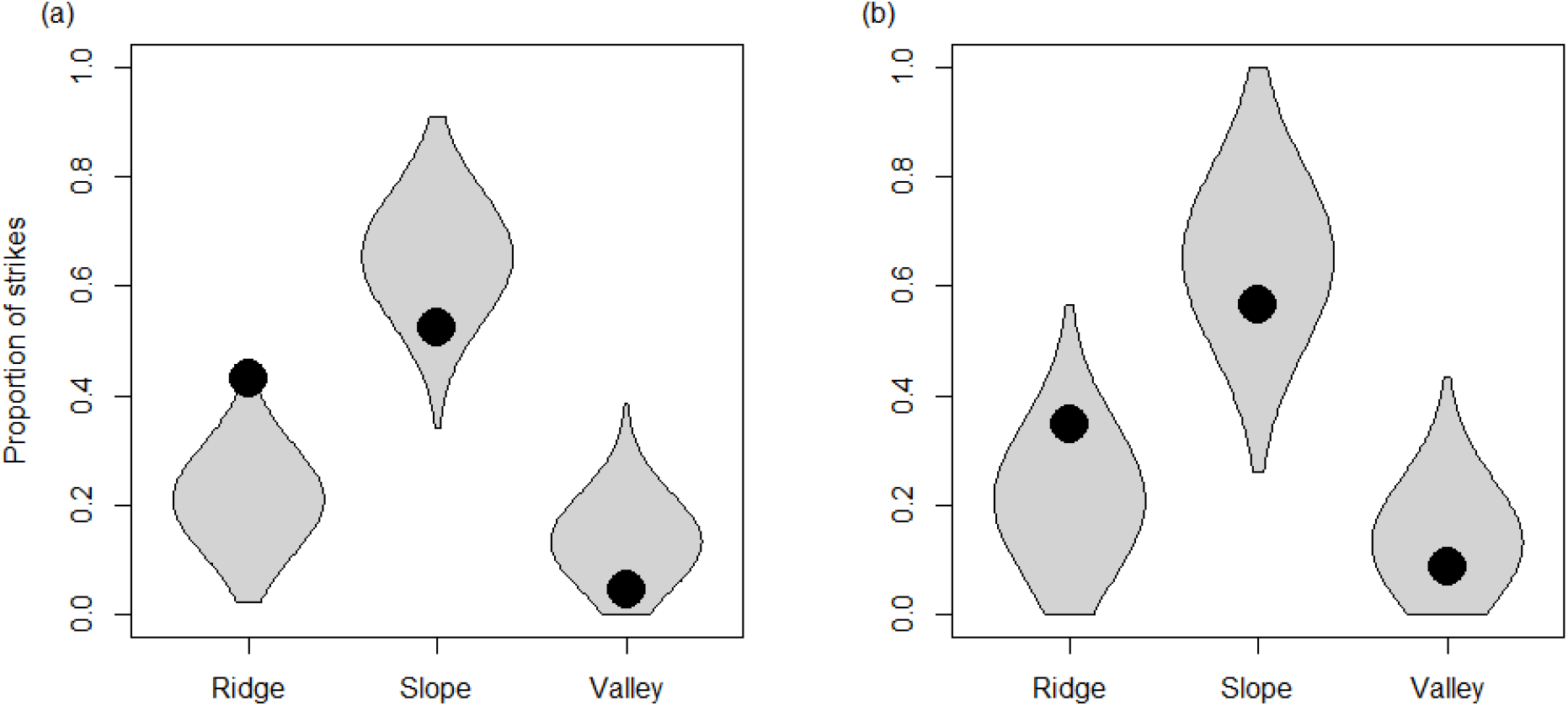
As Figure 1, but restricted to strikes within 10m of the trail (n=25). Across all strikes, they were more likely to occur in ridges (P=0.003) than expected if strikes were randomly distributed with respect to topography, and less likely to occur in slopes (P=0.024) and valleys, although the latter was not significant (P=0.316). Patterns were similar but non-significant for all topography classes for the 15 confirmed strikes (ridges: P=0.091, slopes: P=0.080, valleys: P=0.784).

**Figure S2.**
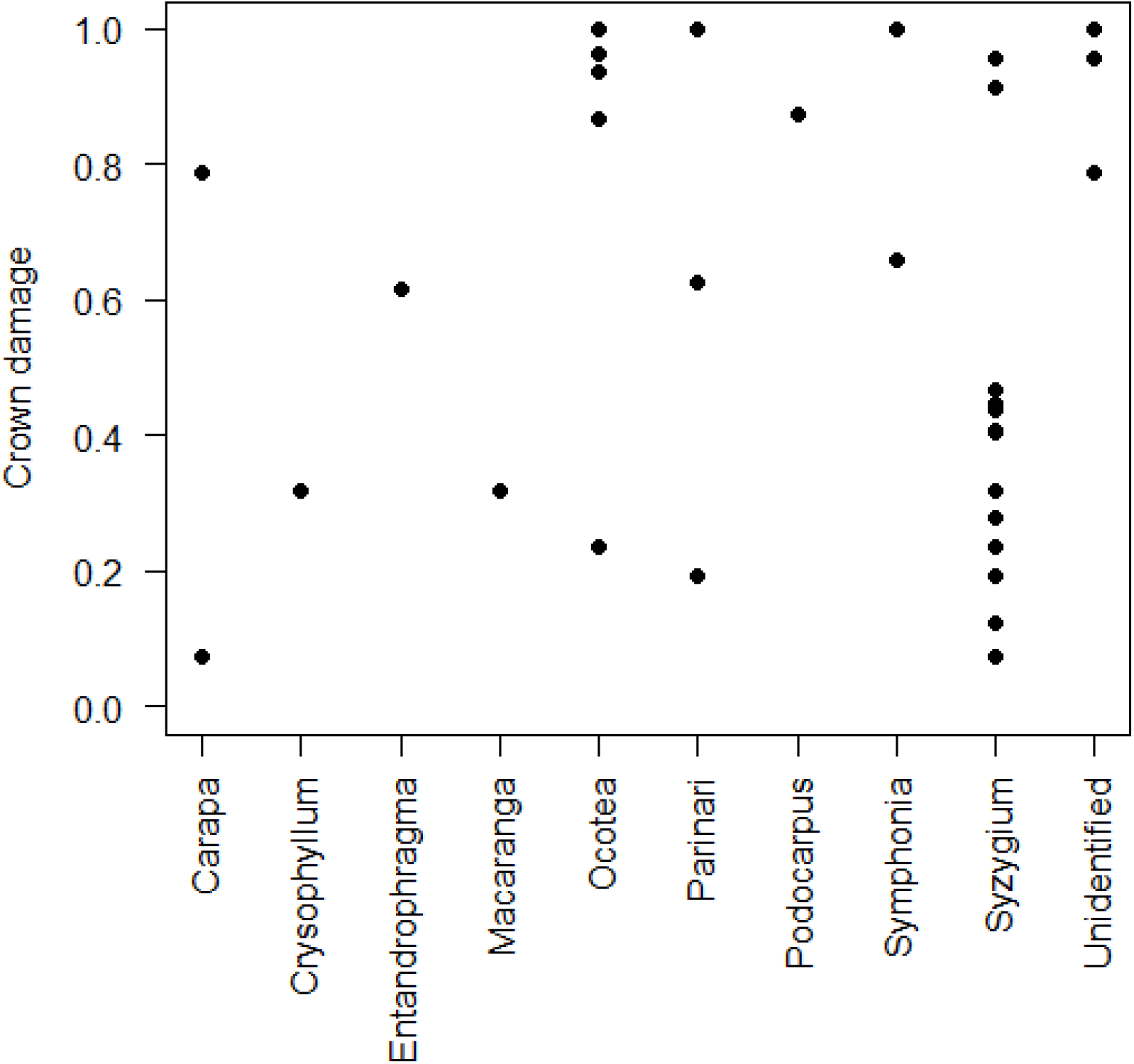
Variation in crown damage between genera. Each point shows crown damage for a tree. All large trees (>50cm DBH) within lightning disturbances have been included.

**Table S1.** Coefficients of generalized liner models relating the number of trees killed or damaged by lightning disturbances, and the proportion of damaged trees that were killed, to topography and forest type.

| Term | Estimate | SE | Z | P |
| --- | --- | --- | --- | --- |
| Number dead |  |  |  |  |
| Intercept (valley, closed canopy) | 0.767 | 0.263 | 2.9 | 0.004 |
| Slope | 0.135 | 0.333 | 0.4 | 0.686 |
| Valley | 0.611 | 0.690 | 0.9 | 0.376 |
| Open forest | -0.649 | 0.323 | -2.0 | 0.044 |
| Number damaged |  |  |  |  |
| Intercept (valley, closed canopy) | 2.468 | 0.147 | 16.8 | <0.001 |
| Slope | 0.126 | 0.181 | 0.694 | 0.487 |
| Valley | -0.027 | 0.427 | -0.1 | 0.950 |
| Open forest | -0.482 | 0.176 | -2.7 | 0.006 |
| Proportion dead |  |  |  |  |
| Intercept (valley, closed canopy) | -1.508 | 0.211 | -7.1 | <0.001 |
| Slope | -0.002 | 0.265 | 0.0 | 0.995 |
| Valley | 0.794 | 0.531 | 1.5 | 0.135 |
| Open forest | -0.190 | 0.261 | -0.7 | 0.466 |
| Number dead - high confidence disturbances |  |  |  |  |
| Intercept (valley, closed canopy) | 1.011 | 0.366 | 2.7 | 0.006 |
| Slope | 0.003 | 0.482 | 0.0 | 0.995 |
| Valley | 0.290 | 0.777 | 0.374 | 0.709 |
| Open forest | -0.494 | 0.454 | -1.1 | 0.277 |
| Number damaged – high confidence disturbances |  |  |  |  |
| Intercept (valley, closed canopy) | 2.82 | 0.184 | 15.3 | <0.001 |
| Slope | -0.035 | 0.241 | -0.1 | 0.884 |
| Valley | -0.417 | 0.425 | -1.0 | 0.326 |
| Open forest | -0.388 | 0.227 | -1.7 | 0.087 |
| Proportion dead – high confidence disturbances |  |  |  |  |
| Intercept (valley, closed canopy) | -1.6 | 0.264 | -6.2 | <0.001 |
| Slope | 0.053 | 0.329 | 0.2 | 0.872 |
| Valley | 0.902 | 0.554 | 1.6 | 0.103 |
| Open forest | -0.115 | 0.3200 | 0.4 | 0.718 |

**Table S2.** Coefficients of linear mixed effects models relating crown damage to taxonomy, topography and forest type.

| Term | Estimate | 95% LCL | 95% UCL | P value |
| --- | --- | --- | --- | --- |
| All strikes |  |  |  |  |
| Intercept | 1.00 | -0.15 | 2.23 | 0.102 |
| Species – Syzygium | -2.03 | -3.26 | -0.83 | <0.001 |
| Topography – Slope | 0.85 | -0.57 | 2.15 | 0.232 |
| Topography – Valley | 1.84 | -0.39 | 3.89 | 0.084 |
| Forest type – Open forest | -0.02 | -1.41 | 1.35 | 0.974 |
| All strikes, no unidentified trees |  |  |  |  |
| Intercept | 0.37 | -0.91 | 1.63 | 0.570 |
| Species – Syzygium | -1.53 | -2.81 | -0.29 | 0.012 |
| Topography – Slope | 0.52 | -0.83 | 2.01 | 0.448 |
| Topography – Valley | 2.20 | -0.11 | 4.50 | 0.066 |
| Forest type – Open forest | 0.52 | -0.91 | 2.01 | 0.474 |
| High-confidence strikes |  |  |  |  |
| Intercept | 0.53 | -0.82 | 1.94 | 0.398 |
| Species – Syzygium | -1.83 | -3.18 | -0.42 | 0.004 |
| Topography – Slope | -0.16 | -1.79 | 1.39 | 0.846 |
| Topography – Valley | 2.32 | 0.26 | 4.34 | 0.026 |
| Forest type – Open forest | 1.75 | 0.16 | 3.49 | 0.018 |
| High-confidence strikes – no unidentified trees |  |  |  |  |
| Intercept | 0.17 | -1.16 | 1.66 | 0.784 |
| Species – Syzygium | -1.47 | -2.95 | -0.22 | 0.030 |
| Topography – Slope | -0.86 | -2.64 | 0.84 | 0.316 |
| Topography – Valley | 2.44 | 0.25 | 4.53 | 0.026 |
| Forest type – Open forest | 2.37 | 0.80 | 4.26 | 0.004 |

